# Comparison of protein and peptide fractionation approaches in protein identification and quantification from *Saccharomyces cerevisiae*

**DOI:** 10.1101/2020.02.13.948513

**Authors:** Liting Deng, David C. L. Handler, Dylan Multari, Paul A. Haynes

**Affiliations:** Department of Molecular Sciences, Faculty of Science and Engineering, Macquarie University, Sydney, NSW, Australia

**Keywords:** proteome, yeast, SDS-PAGE, FASP, GPF, high pH reversed phase fractionation

## Abstract

Proteomics, as a high-throughput technology, has been developed with the aim of investigating the maximum number of proteins in cells. However, protein discovery and data generation vary in depth and coverage when different technical strategies are used. In this study, four different sample preparation, and peptide or protein fractionation, methods were applied to identify and quantify proteins from log-phase yeast lysate: sodium dodecyl sulfate-polyacrylamide gel electrophoresis (SDS-PAGE); gas phase fractionation (GPF); filter-aided sample preparation (FASP)- GPF; and FASP-high pH reversed phase fractionation (HpH). Fractionated samples were initially analyzed and compared using nanoflow liquid chromatography-tandem mass spectrometry (LC-MS/MS) employing data dependent acquisition on a linear ion trap instrument. The number of fractions and replicates was adjusted so that each experiment used a similar amount of mass spectrometric instrument time, approximately 16 hours. A second set of experiments was performed using a Q Exactive Orbitrap instrument, comparing FASP-GPF, SDS-PAGE and FASP-HpH. Compared with results from the linear ion trap mass spectrometer, the use of a Q Exactive Orbitrap mass spectrometer enabled a small increase in protein identifications using SDS-PAGE and FASP-GPF methods, and a large increase using FASP-HpH. A big advantage of using the higher resolution instrument found in this study was the substantially increased peptide identifications which enhance the proteome coverage. A total of 1035, 1357 and 2134 proteins were separately identified by FASP-GPF, SDS-PAGE and FASP-HpH. Combining results from the Orbitrap experiments, there were a total of 2269 proteins found, with 94% of them identified using the FASP-HpH method. Therefore, the FASP-HpH method is the optimal choice among these approaches when using a high resolution spectrometer, when applied to this type of sample.

## 1. INTRODUCTION

Proteins are essential components of the cellular machinery, performing and enabling precise tasks in highly complex living biological systems. Life depends on these versatile macromolecules and their ability to perform complex biological roles from genetic replication to cell senescence and death [1, 2]. Bottom-up shotgun proteomics based on mass spectrometry is an extremely powerful tool to use in proteome analysis since it enables the identification and characterization of peptides, proteins and their modifications [3, 4]. The general procedure of proteomic analysis includes sample preparation, mass spectrometry (MS) data acquisition, and statistical and bioinformatics analysis [1, 5, 6].

A number of techniques and strategies have been developed in recent years, which are part of the ongoing improvements in the entire process of proteomics analysis. However, one of the limits in current proteomic technologies is that there is no single standardized strategy allowing for the analysis of the entire proteome in a simple step. The large number of peptides and the wide dynamic range of protein concentration in the proteome of different biological samples make the proteomic analysis a complex problem [7]. In large-scale proteomic studies, mass spectrometry results and the depth of coverage of the proteome have remained largely depend on the sample preparation, which makes it the most crucial part in the whole proteomics process [8]. Additionally, optimal sample preparation methods are necessary in order to obtain reliable results, particularly in comparative proteomics where the identification of differentially expressed proteins can be heavily influenced by even minor differences in sample preparation [9].

Sample preparation should ideally be as simple as possible in order to reduce time and avoid sample loss, while eliminating interferences or contaminants [10]. Theoretically, the best strategy for sample preparation would be no sample preparation at all, but with effective protein identification and maximal depth of coverage [11]. For example, a simple non-fractionated approach may also give comparable results while without requiring extra machine-time and analysis [11]. However, the complexity of the typical dynamic proteome far exceeds the capacity of currently available analytical systems, so some pre-fractionation is generally necessary. Notwithstanding the ongoing development of new technologies, the initial step in any proteomic experiment is to reduce the sample complexity by carefully preparing and fractionating the samples at either the protein or peptide level with the aim of obtaining a more comprehensive data profile. Commonly used approaches include techniques such as SDS-PAGE for separation of proteins based on their molecular weights, gas phase fractionation (GPF) of peptides based on the mass of their ions detected in the mass spectrometer, and high pH reversed phase fractionation (HpH) for peptide separation by hydrophobicity [12, 13].

The in-gel protein digestion protocol for proteomics, established over 20 years ago, has been the cornerstone method affording robust protein identifications for many ongoing proteome projects [14, 15]. Gel bands are manually cut and fractionated according to the molecular weights of separated proteins. Peptides are recovered from the gel slices after in-gel trypsin digestion, and then introduced into a mass spectrometer for analysis, typically by reversed phase peptide separation [16]. Preparing more gel fractions decreases the complexity of the individual samples, which can have a substantial impact on the depth of proteome analysis [17]. It has been reported previously that a combination of in-solution digestion and peptide fractionation methods could provide higher peptide recovery than the protein fractionation method based on SDS-PAGE [18, 19]. Contrasting findings have been reported in other studies, which serves to highlight that such conclusions are strongly sample dependent [20, 21].

The GPF method utilizes the resolving power of the mass spectrometer to separate ions of different mass ranges to greatly reduce the complexity of peptide mixtures [22]. Peptides are repetitively injected into the mass spectrometer, and only a relatively narrow m/z range is interrogated by data-dependent precursor ion selection in each analysis. Compared to the ions selected from the wide mass range scan in typical LC-MS/MS analysis, a smaller m/z range in each GPF experiment is set for precursor ion selection, so that the components within different mass ranges can be better resolved to reduce the complexity of analysis [18, 22]. GPF has been described as a means to achieve higher proteome coverage than multiple LC-MS/MS analysis of unfractionated complex peptide mixtures [17, 23]. A combination of in-solution trypsin protein digestion with GPF has been shown to reduce sample preparation time, and increase the number of proteins identified [17, 18, 24]. However, the main concern of GPF is that only a small portion of the whole sample is actually analyzed by the mass spectrometer in each analysis, which may lead to loss of information [20].

Another method to reduce the complexity of peptide mixture is HpH fractionation. Peptides are bound to the hydrophobic resin under aqueous conditions and desalted by washing the column with water. A step gradient of increasing ACN concentrations in a volatile high pH elution solution is then applied to the columns to elute bound peptides into different fractions which can be collected by centrifugation. During MS analysis, peptides in each high pH fraction are further separated using a low-pH reversed phase gradient which separates peptides via an orthogonal mechanism, thus reducing the overall sample complexity and improving the ability to identify low abundant peptides [25, 26].

FASP has become a widely used method for in-solution digestion of proteins due to its ability to remove detergents prior to mass spectrometry analysis [5]. The FASP method uses a common ultrafiltration device whereby the membrane pores are small enough to allow contaminating detergents to pass through, while proteins are retained and concentrated in the filter unit [24]. The combination of FASP based in-solution digestion and GPF has been shown previously to be highly effective in achieving high-quality proteome coverage [19].

In this study, four proteomics analysis strategies (SDS-PAGE, GPF, FASP-GPF and FASP-HpH) were compared. Aliquots of 100 μg of proteins extracted from yeast BY4741 cells were used for each experiment. In the first stage of the study, SDS-PAGE, GPF, FASP-GPF and FASP-HpH were compared in terms of protein and peptide identification using a linear ion trap mass spectrometer. For SDS-PAGE fractionations, replicate gels were separately cut into 8, 16 and 32 slices to investigate the impact of the number of fractions analyzed on protein identifications. Similarly, separate 100 μg protein aliquots were digested in solution in both Eppendorf tubes and FASP filters. Replicate analyses were performed with GPF using 4, 8 and 16 mass range windows, to investigate the impact of mass range selection and replicate number on protein identifications. Considering both analysis time required and number of proteins identified, SDS-PAGE (16 fractions), FASP-GPF (4 mass ranges, 4 replicates) and FASP-HpH (8 fractions, 2 replicates) methods were found to be higher performing strategies. These experiments were replicated using a high resolution Orbitrap mass spectrometer in the second stage of the study, with an equal amount of instrument time used for each of the approaches examined. Proteins identified were categorized in terms of abundance by correlation with published mRNA data, sorted by known subcellular localization, and classified according to protein function using available gene ontology information. Results from all these experiments and analyses were quantitatively compared in order to determine the optimal method for proteome analysis with regard to this particular type of sample.

## 2. MATERIALS AND METHODS

### 2.1 Cell culture

*Saccharomyces cerevisiae* BY4741 single colonies were selected from a YPD plate (1% Yeast extract, 2% peptone, 2% dextrose, and 2% agar) using sterile pipette tips. Cells were separately dissociated into three tubes as biological replicates, with 10 mL YPD medium (1% Yeast extract, 2% peptone, and 2% dextrose) by pipetting up and down. Samples were cultured overnight on a shaking incubator (30°C, 200 rpm). 100 μL cells were transferred and cultured in 250 mL flask with 100 mL YPD medium. Cells were harvested prior to stationary phase by centrifugation at 4℃ (2500 rpm, 10 min) and then washed twice with pre-cooled Milli-Q water. Samples were stored at −20℃ until protein extraction.

Three biological replicate cell cultures were grown and harvested for each subsequent experiment involving different sample preparation and fractionation method comparisons.

### 2.2 Protein extraction

Yeast cells were pre-treated with 2 M lithium acetate, and then with 0.4 M NaOH for 5 min on ice. Cell pellets were re-suspended with 100 μL SDS-PAGE sample buffer and heated to 100℃ for 5 min. Lysates were centrifuged at 13,000 rpm for 10 min to clear cellular debris [27]. The supernatant was carefully transferred to a fresh tube and then precipitated to remove interfering agents using methanol/chloroform (sample: methanol: chloroform: water = 1:4:1:3) [28]. Protein pellets were dissolved in 8 M urea/100 mM Tris-HCl buffer (pH 8.0) and the concentration was quantified with bicinchoninic acid (BCA) protein assay kit (Pierce, Rockford, LI, USA) with bovine serum albumin as standards.

### 2.3 In-gel digestion

Aliquots containing 100 μg of protein were mixed with 2× SDS sample buffer and heated at 95°C for 10 min. The mixture was loaded onto Bio-Rad 10% TGX mini gels and separated at 120 V for 65 min. Gels were stained with colloidal Coomassie blue G-250 for 40 min and then de-stained with Milli-Q water overnight. Individual gel lanes from each biological sample were separately sliced into 8, 16 and 32 equal sized pieces and finely chopped, with three replicates performed of each fractionation regime. Gel pieces were washed once with 100 mM NH_4_HCO_3_, and twice with 50% ACN/50 mM NH_4_HCO_3_ (10 min), and then dehydrated with 100% ACN (5 min). Samples were air-dried and reduced with 50 μL of 10 mM DTT in 100 mM NH_4_HCO_3_ for 45 min at room temperature. DTT was removed and then followed with alkylation in dark for 45 min by adding 50 μL of 55 mM iodoacetamide in 100 mM NH_4_HCO_3_. After alkylation, gels were briefly washed once with 100 mM NH_4_HCO_3_ (5 min), twice with 50% ACN/50 mM NH_4_HCO_3_ (5 min), then dehydrated with 100% ACN (5 min). Samples were air-dried, trypsin was added to the sample for equilibration at 4°C for 60 min before digestion at 37°C over-night. Peptides were extracted from gel pieces using 50% ACN/2% formic acid for 30 min. The peptide extraction was repeated twice. Tryptic peptides were combined and evaporated to dryness in a vacuum centrifuge and reconstituted with 10 μL of 1% formic acid.

### 2.4 In-solution digestion

Protein samples were resuspended in 8M urea buffer and diluted with 100 mM Tris-HCl (pH 8.0) to a final urea concentration of 1.6 M before adding trypsin and incubating at 37°C over-night. The pH was adjusted by formic acid to 2~3 before peptides were purified using the SDB-RPS (3M-Empore) stage-tips. Peptides were eluted twice with 200 μL 80% ACN/5% ammonium hydroxide. Extracts were combined and evaporated to dryness in a vacuum centrifuge and reconstituted with 160 μL of 1% formic acid for MS analysis.

### 2.5 FASP-based in solution digestion

Proteins were reduced in 200 μL of 50% Trifluoroethanol (TFE)/100 mM NH_4_HCO_3_/50 mM DTT for 60 min, and then concentrated to 20 μL in Amicon Ultra 0.5 mL ultrafiltration devices (10 K cut-off, Millipore) by centrifugation at 13,000 rpm for 30 min. 200 μL of 50% TFE/100 mM NH_4_HCO_3_ /60 mM iodoacetamide was added to the mixture and proteins were alkylated in the dark for 45 min. Reagents were removed by centrifugation at 13,000 rpm for 30 min. Concentrated proteins were washed three times with addition of 200 μL of 50% TFE/100 μM NH_4_HCO_3_ and centrifugation at 13,000 rpm for 30 min. Proteins were digested in 300 μL of 20% ACN/100 mM NH_4_HCO_3_ buffer with 2 μg trypsin (Promega) and incubated at 37 °C overnight in the ultrafiltration device. Tryptic peptides were centrifuged into new ultrafiltration receptacles. This process was followed by another two rinses using 150 μL of 50% ACN/2% formic acid. Extracts were combined and evaporated to dryness in a vacuum centrifuge and re-constituted with 160 μL of 1% formic acid.

### 2.6 High pH reversed phase fractionation

Aliquots of 100 μg of tryptic peptides were resuspended in 300 μL of 0.1% Trifluoroacetic acid and loaded to the pre-conditioned spin column from the high pH reversed-phase peptide fractionation kit (Thermo Fisher, CA, USA). After sample was washed once with water by low speed centrifugation (3000 g, 2 min), a step gradient of increasing ACN concentrations in triethylamine (5%, 7.5%, 10%, 12.5%, 15%, 17.5%, 20% and 50%) was applied to the column to elute bound peptides into eight different fractions by centrifugation (3000 rpm, 2min). Each fraction was then dried in a vacuum centrifuge and reconstituted with 20 μL of 1% formic acid, allowing for each fraction to be analyzed in duplicate using 10 μL aliquots.

### 2.7 Analysis of tryptic digests using SDS-PAGE and FASP-HpH methods on a linear ion trap mass spectrometer

Each sample from in-gel tryptic digestion and high pH reversed phase fractionation were analyzed on a Velos pro linear ion trap mass spectrometer connected to an Easy-nLC 1000 nanoflow HPLC system (Thermo, CA, USA). Chromatography was performed on a 100 um I.D reversed phase column packed in-house to 10 cm with Aqua C18 beads (200 Å, 3 μm) in a fused silica capillary with an integrated electrospray tip which was coupled with a 3 cm pre-column packed with the same. A 1.8 kV electrospray voltage was applied via a liquid junction upstream of the analytical column. Peptides were loaded in buffer A (2% v/v ACN, 0.1% v/v formic acid) and a gradient was developed using buffer B (99.9% v/v ACN, 0.1% v/v formic acid) as follows: 0-30% buffer B at flow 550 nL/min (36min), 30%−50% buffer B at flow 600 nL/min (12min), 50%−95% buffer B at flow 600 nL/min (2 min), 95% buffer B at flow 850 nL/min (10 min). Spectral acquisition was performed in positive ion mode over the scan range of 400 m/z to 1500 m/z using Xcalibur software (Thermo, v2.07). A normalized collision energy of 35% was used to perform MS/MS of the top nine most intense precursor ions, with dynamic exclusion enabled for 90 s [29–31].

### 2.8 Analysis of in-solution tryptic digests using GPF on a linear ion trap mass spectrometer

Mass ranges for GPF windows were calculated according to the frequency distribution of peptides, based on preliminary experiments in which 10 μg of yeast lysate peptides was loaded onto the nanoLC without pre-fractionation and separated using a 180 min gradient (data not shown). In subsequent experiments, 4, 8, and 16 mass ranges were separately set as follows: four mass ranges (400–665, 660–835, 830–1005 and 1000–1500 amu), eight mass ranges (400-585, 580-665, 660-745, 740-835, 830-905, 900-1005, 1000-1155, 1150-1500 amu) and 16 mass ranges (400-515, 510-585, 580-625,620-665, 660-715, 710-745, 740-795, 790-835, 830-875, 870-905, 900-955, 950-1005, 1000-1075, 1070-1155, 1100-1265, 1260-1500 amu). There were 16 injections (10 μL was loaded each time) applied for each of three sample replicates as mentioned previously. Other settings were as the same as the method above which was used for analyzing the in-gel digested peptides.

### 2.9 Analysis of tryptic digests on a Q Exactive Orbitrap mass spectrometer

Trypsin digests from experiments using FASP-GPF (4 mass ranges, 4 replicates), SDS-PAGE (16 fractions), and FASP-HpH (8 fractions, 2 replicates) were analyzed on a Q Exactive Orbitrap mass spectrometer (Thermo Scientific, San Jose, CA, USA) coupled to an EASY-nLC1000 nanoflow HPLC system (Thermo Scientific, San Jose, CA, USA). Reversed-phase chromatographic separation was performed on an in-house packed reverse–phase column (75 μm × 10 cm with Halo 2.7 μm 160 Å ES-C18 beads (Advanced Materials Technology). Peptides were separated for 60 min using a gradient of solvent A (97.9% water/2% acetonitrile/0.1% formic acid) and solvent B (99.9% acetonitrile/0.1% formic acid) at a flow rate 300 nL/min as follows: 1-50% buffer B (50 min), 50%-85% buffer B (2 min), 85% buffer B (8 min). To identify proteins, full MS scan spectra from m/z 350 ~2000 amu were acquired at resolution of 35,000 and an automatic gain control target value of 1 × e^6^ ions. For FASP-GPF (4 mass ranges, 4 replicates) method, the same mass ranges setting were kept, to enable comparisons with previous experiments using the same method on the ion trap mass spectrometer. The top 10 most abundant ions were selected with precursor isolation width of 3 m/z for higher-energy collisional dissociation (HCD) fragmentation. HCD normalized collision energy was set to 30% and fragmented ions were detected in the Orbitrap at a resolution of 17,500 with an automatic gain control target value of 2 × e^5^ ions. Target ions that had been selected for MS/MS were dynamically excluded for 20 s.

### 2.10 Peptide to spectrum matching

Raw files from all experiments were converted to mzXML format using readWbatch, and peptide to spectrum matching was performed using the X! Tandem algorithm [32, 33]. Fractions were processed sequentially and then merged for each sample replicate. Non-redundant output files were generated for protein identifications with log (e) values less than −1. MS/MS spectra were searched against the *S. cerevisiae* protein sequence database (6800 sequence entries, ENSEMBL), as well as common human and trypsin peptide contaminants, and additional searching was performed against a reversed sequence database to evaluate the FDR. Search parameters included MS and MS/MS tolerances of ± 4 Da and ± 0.4 Da, respectively. The criteria for the database search included carbamidomethylation (C) as complete modification, acetylation (N) and oxidation (M) as potential modification, with fully tryptic cleavage sites.

### 2.11 Statistical and bioinformatic analysis

Low stringency search data from peptide to spectrum matching outputs from X! Tandem for individual replicates was transformed into high stringency data by combining results from three biological replicate experiments into a single list of reproducibly identified proteins using Scrappy [34] and PeptideWitch [35]. The criteria for a reproducibly identified protein was that a protein must be present in all three replicates, with a minimum peptide spectral count of five [36–38]. Protein FDR in reproducibly identified proteins from all experiments in this study was no more than 2%, and peptide FDR was less than 0.3%, indicating that no further filtering was required. Reproducibly identified proteins from different experiments were classified according to their known function based on gene ontology information in the Saccharomyces Genome Database using the GO Slim mapper function [39, 40]. This function was also performed for localization analysis of identified proteins based on the cellular component ontology information.

The correlation of mRNA expression and protein abundance was analyzed using PARE [41]. Proteins identified from different experiments were analyzed to compare the performance of each of these methods in identifying proteins of varying abundance, based on information from available mRNA databases. Messenger RNA data of yeast BY4741, also cultured in YPD medium and harvested during log phase, was downloaded from NCBI with GEO series accession number GSE9217 and set as a reference for the proteins identified in this study (5717 identifications from three replicates) [42]. Proteins identified in specific experiments were firstly matched to identifications from the mRNA reference. The matched proteins and their expression, and the mRNA reference, were uploaded to PARE, and a good correlation between mRNA abundance and protein expression was found. mRNAs with higher abundance corresponded to more protein expression, therefore, the abundance of mRNA from the reference was used to indicate the theoretical value of the corresponding protein expression. mRNA identifications were ranked based on their normalized abundance by PARE and distributed into six categories, and these corresponding proteins in specific groups were counted, and the percentage of these proteins in total matched proteins to the mRNA reference in specific experiments were calculated and compared.

## 3. RESULTS

### 3.1 Proteins identified with SDS-PAGE fractionation on a linear ion trap mass spectrometer

The number of reproducibly identified proteins and peptides identified using SDS-PAGE with different numbers of fractions is presented in Table 1, along with a brief summary of reproducibly identified proteins in all experiments performed in this study. A total of 763, 1206, and 1329 proteins were reproducibly identified from yeast samples with 8, 16, and 32 fractions, respectively. Of these, 718 proteins were found in all three experiments, which indicates a high degree of experimental consistency (Figure 1A). As expected, more proteins and peptides were detected with an increase in the number of gel fractions. There were 60% more proteins identified with 16 fractions than 8 fractions, but only 10% more proteins identified with 32 fractions when compared with 16 fractions. This shows that using eight fractions does not simplify the complex peptide mixtures sufficiently, because expanding to 16 fractions produces a large increase in protein identifications. On the other hand, moving from 16 fractions to 32 fractions produced a much smaller increase in protein identifications, suggesting that in the 16 fractions experiment the complexity of the peptide mixture is compatible with the analytical capacity of the instrument. The advantage gained by identifying slightly more proteins is outweighed by the disadvantage of requiring twice as much instrument time to complete the analysis.

**Table 1.** A brief summary of identified proteins in each experiment separately applied on linear ion trap and Q Exactive Orbitrap mass spectrometers.

| Instruments | Methods | No. of fractions or mass ranges | Analysis repeats per fraction or mass range | No. of injections | Gradient per injection (hrs) | Analysis time per sample (hrs) | No. of reproducibly identified proteins | No. of reproducibly identified peptides | Protein FDR | Peptide FDR |
| --- | --- | --- | --- | --- | --- | --- | --- | --- | --- | --- |
| Linear ion trap | **SDS-PAGE** | 8 | 1 | 8 | 1 | 8 | 763 | 16199 | 0.13% | 0.03% |
| Linear ion trap | **SDS-PAGE** | 16 | 1 | 16 | 1 | 16 | 1206 | 40599 | 1.78% | 0.28% |
| Linear ion trap | **SDS-PAGE** | 32 | 1 | 32 | 1 | 32 | 1329 | 65006 | 2.00% | 0.26% |
| Linear ion trap | **GPF** | 4 | 4 | 16 | 1 | 16 | 642 | 17815 | 1.68% | 0.30% |
| Linear ion trap | **FASP-GPF** | 4 | 4 | 16 | 1 | 16 | 698 | 15863 | 0.57% | 0.18% |
| Linear ion trap | **FASP-GPF** | 8 | 2 | 16 | 1 | 16 | 479 | 8485 | 0.62% | 0.14% |
| Linear ion trap | **FASP-GPF** | 16 | 1 | 16 | 1 | 16 | 415 | 4665 | 0.00% | 0.00% |
| Linear ion trap | **FASP-HpH** | 8 | 2 | 16 | 1 | 16 | 683 | 17588 | 2.00% | 0.41% |
| Q Exactive | **FASP-GPF** | 4 | 4 | 16 | 1 | 16 | 1035 | 29062 | 1.98% | 1.14% |
| Q Exactive | **SDS-PAGE** | 16 | 1 | 16 | 1 | 16 | 1357 | 112052 | 1.72% | 0.26% |
| Q Exactive | **FASP-HpH** | 8 | 2 | 16 | 1 | 16 | 2134 | 167337 | 1.52% | 0.19% |

**Figure 1.**
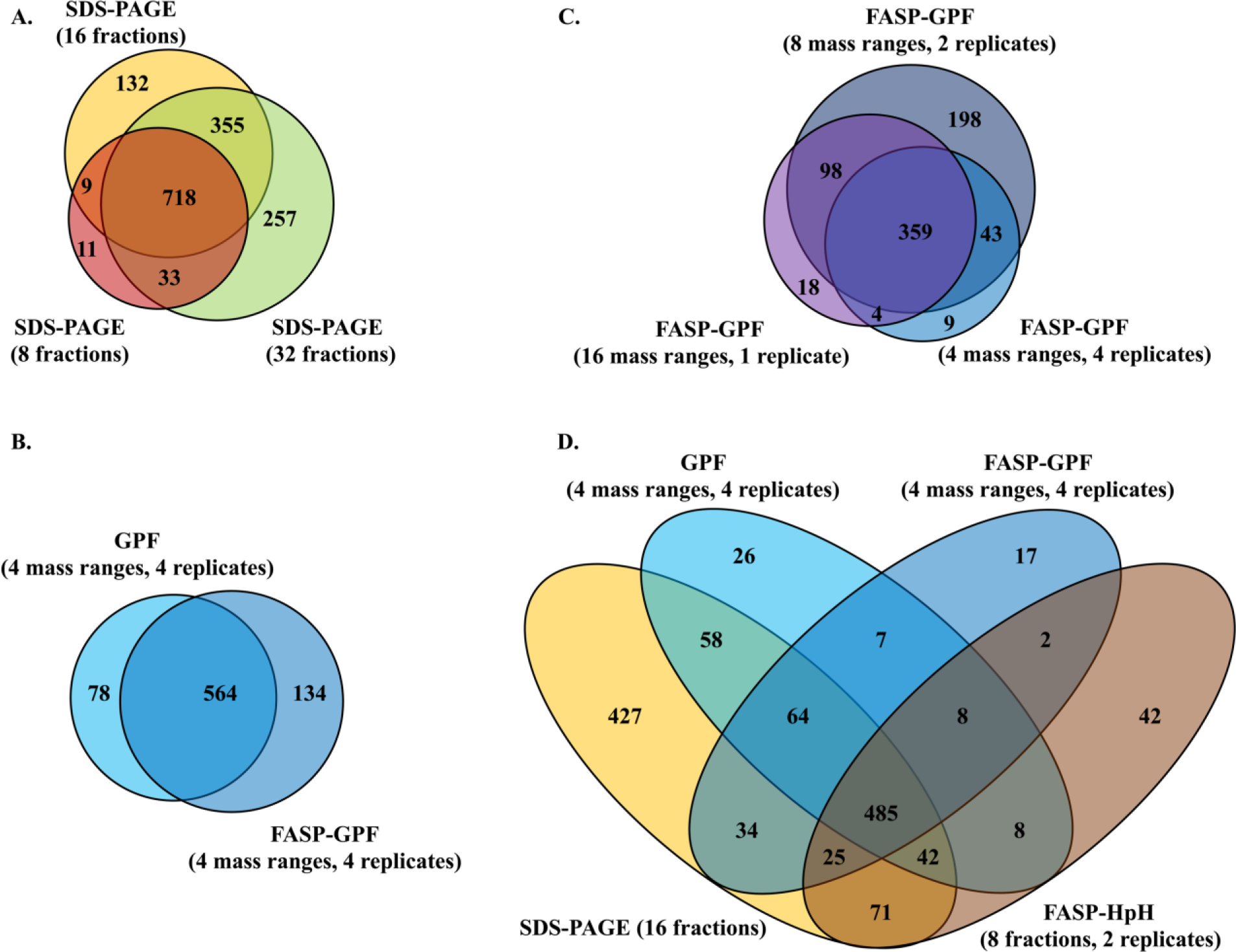
Venn diagram analysis of proteins identified using different methods on linear ion trap mass spectrometer. (A) Venn diagrams analysis of proteins separately identified using SDS-PAGE (8 fractions), SDS-PAGE (16 fractions), and SDS-PAGE (32 fractions). (B) Venn diagram analysis of proteins separately identified using GPF (4 mass ranges, 4 replicates) and FASP-GPF (4 mass ranges, 4 replicates). (C) Venn diagram analysis of proteins separately identified using FASP-GPF (4 mass ranges, 4 replicates), FASP-GPF (8 mass ranges, 2 replicates), and FASP-GPF (16 mass ranges, 1 replicate). (D) Venn diagrams analysis of proteins separately identified using SDS-PAGE (16 fractions), GPF (4 mass ranges, 4 replicates), FASP-GPF (4 mass ranges, 4 replicates) and FASP-HpH (8 fractions, 2 replicates).

### 3.2 Proteins identified with GPF method on a linear ion trap mass spectrometer

In GPF experiments, a sample was repeatedly analyzed by automated data-dependent tandem MS using multiple narrow m/z ranges for survey scans from which to select ions for fragmentation, rather than a single wide range survey scan. As shown in Table 1, the use of four mass range windows repeated four times gave the best results, with 642 proteins identified in the solution digested GPF experiment and 698 proteins identified in the FASP-GPF experiment, with 564 proteins found in both (Figure 1B). Hence, the FASP-GPF method allowed for the identification of approximately 10% more reproducibly identified proteins than the GPF experiment without using FASP. Moving to a greater number of mass ranges and less replicates, somewhat surprisingly, produced approximately 40% less protein identifications. As shown in Table 1, the use of eight mass windows and two replicates identified 479 proteins, while the use of 16 mass windows in a single replicate experiment reduced 283 protein identifications. A total of 359 proteins were found in all three experiments (Figure 1C).

### 3.3 Comparative analysis of proteins identified from SDS-PAGE, in solution GPF, FASP-GPF and FASP-HpH using a linear ion trap mass spectrometer

As depicted in Table 1, there were 683 proteins and 17588 peptides identified using FASP-HpH. which is comparable with the experimental outputs using GPF methods with four mass ranges, and the SDS-PAGE method with eight fractions. With the consideration of both instrument time and protein identifications, SDS-PAGE (16 fractions), GPF (4 mass ranges, 4 replicates), FASP-GPF (4 mass ranges, 4 replicates) and FASP-HpH (8 fractions, 2 replicates) are concluded as the four best performing experimental approaches with a total of 1316 non-redundant proteins identified. A total of 485 proteins were found to be overlapping between all four as shown in Figure 1D. SDS-PAGE clearly performed better in identifying more proteins compared to the other three methods. The majority of proteins identified by GPF and FASP-HpH experiments were also found using the SDS-PAGE method, while 427 unique proteins were only reproducibly detected in SDS-PAGE based experiments. However, it must be noted that there were some proteins uniquely identified by each technique.

The average count of total peptides per protein identified in each technique (SDS-PAGE, GPF, FASP-GPF and FASP-HpH with 16 injections) was found to be 34 peptides per protein for SDS-PAGE, 28 peptides per protein for in solution digestion and GPF, 23 peptides per protein for FASP-GPF, and 26 peptides per proteins for FASP-HpH. Additionally, the average count of peptides per protein from the 485 overlapping proteins identified in all four experiments was also analyzed. For those particular proteins, 71, 35, 30 and 34 total peptides per protein were discovered using SDS-PAGE, GPF, FASP-GPF and FASP-HpH, respectively. These results show that SDS-PAGE clearly performed better in identifying more peptides per protein, which equates to enhanced proteome coverage. For example, the redundant count of peptides identified from protein TDH3 by SDS-PAGE, GPF, FASP-GPF and FASP-HpH were 1735, 884, 663 and 732, respectively. Even when considering nonredundant accounting of peptides so that each unique peptide is counted only once, the SDS-PAGE approach still performed better, with 79, 56, 61 and 58 unique peptides identified from TDH3 in the four experiments.

The proteins identified using SDS-PAGE, GPF, FASP-GPF and FASP-HpH were sorted using their theoretical molecular weight values (Figures 2). The SDS-PAGE method allowed a greater number of total (Figure 2A) and unique proteins (Figure 2B) to be identified across all different molecular weight ranges, especially from 0-60 kDa. These results indicate that SDS-PAGE is a broadly applicable fractionation technique to identify more proteins and peptides from yeast lysate than the other fractionation methods.

**Figure 2.**
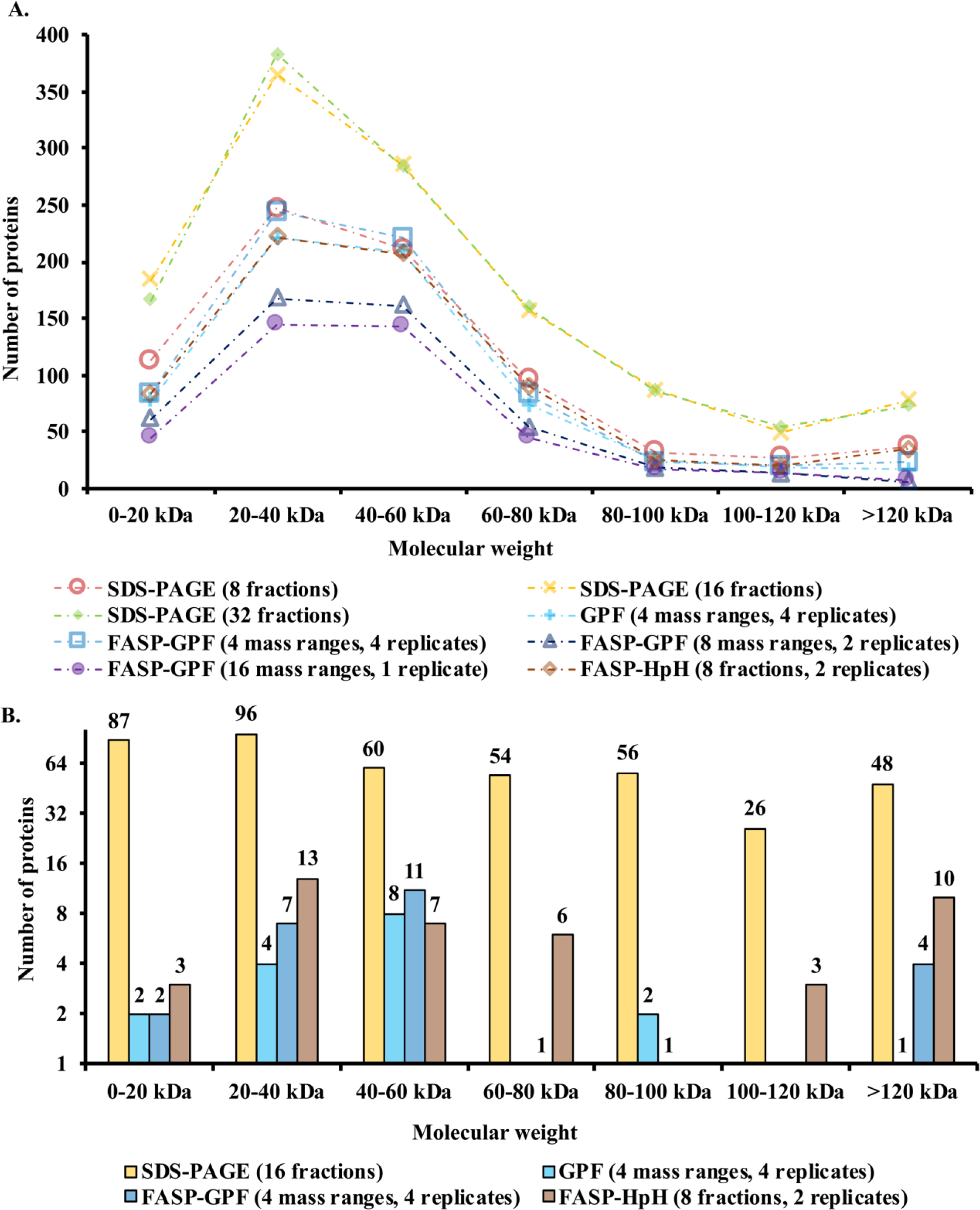
(A) Molecular weight distribution analysis of proteins separately identified using SDS-PAGE (8 fractions), SDS-PAGE (16 fractions), SDS-PAGE (32 fractions), GPF (4 mass ranges, 4 replicates), FASP-GPF (4 mass ranges, 4 replicates), FASP-GPF (8 mass ranges, 2 replicates), FASP-GPF (16 mass ranges,1 replicate), and FASP-HpH (8 fractions, 2 replicates) on linear ion trap mass spectrometer. (B) Molecular weight distribution of unique proteins separately identified using SDS-PAGE (16 fractions), GPF (4 mass ranges, 4 replicates), FASP-GPF (4 mass ranges, 4 replicates) and FASP-HpH (8 fractions, 2 replicates) on linear ion trap mass spectrometer.

### 3.4 Correlation with mRNA abundance of proteins identified from SDS-PAGE, GPF, FASP-GPF and FASP-HpH experiments using a linear ion trap mass spectrometer

The correlation analysis of protein abundance and mRNA expression indicated that there was no significant difference between 16 and 32 fractions using SDS-PAGE methods in identifying proteins with lower abundance, as shown in Figure 3. FASP-GPF with four mass ranges performed better in identifying proteins with lower expression level when compared with 8 and 16 mass ranges, and the FASP-HpH strategy, GPF and FASP-GPF techniques using four mass ranges showed similar performance in identifying less abundant proteins (Figure 3). However, the SDS-PAGE fractionation technique was able to detect more proteins with lower abundance than GPF, FASP-GPF and FASP-HpH.

**Figure 3.**
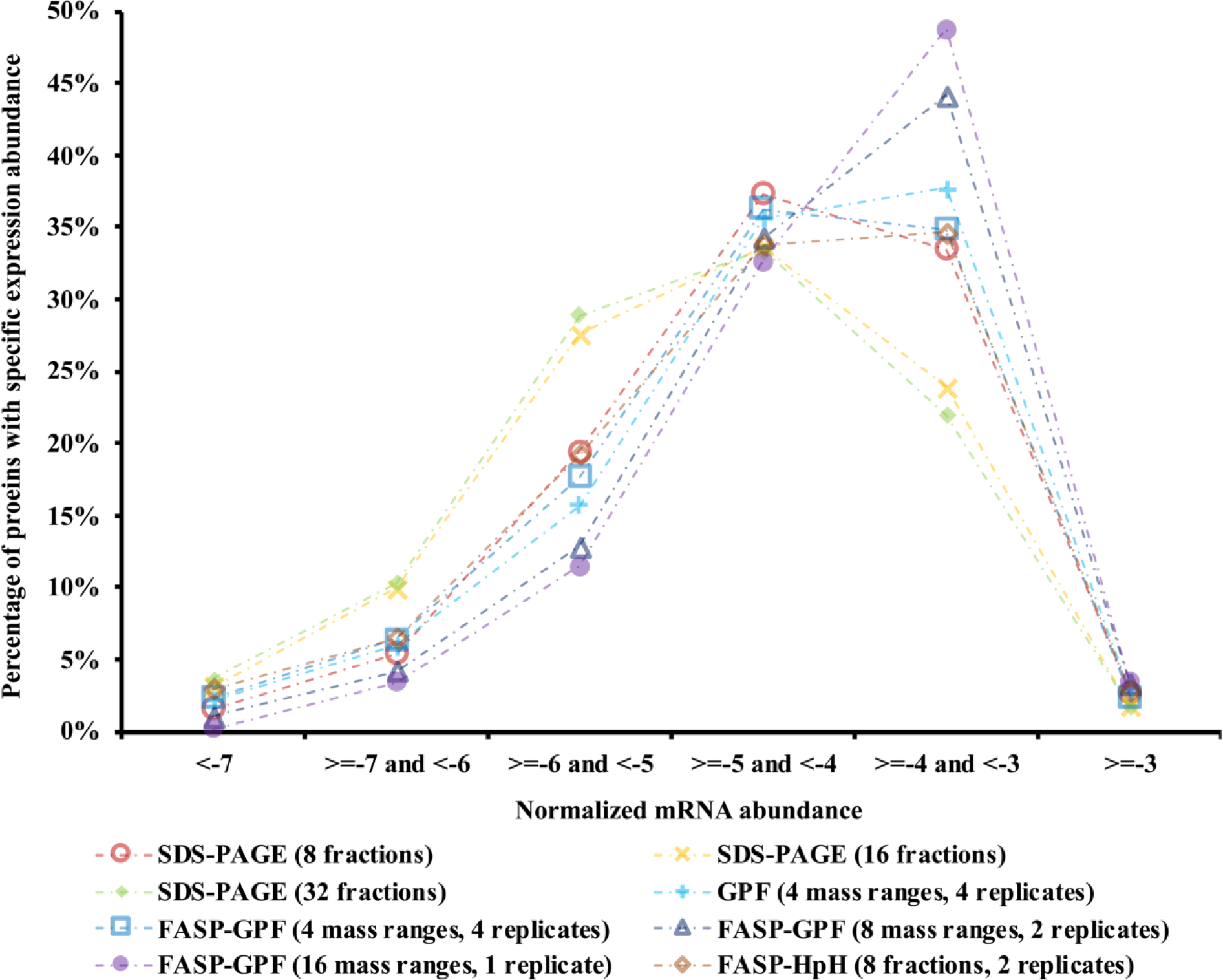
The correlation of protein abundance and mRNA expression was analyzed using PARE, as described in methods. The X axis represents the normalized abundance of mRNAs which were categorized into six groups and used for indicating the theoretical value of corresponding protein expression. The minimum number indicates the category with the lowest abundance. The Y axis represents the percentage of proteins with specific expression abundance in all matched proteins to the mRNA reference in specific experiments, including SDS-PAGE (8 fractions), SDS-PAGE (16 fractions), SDS-PAGE (32 fractions), GPF (4 mass ranges, 4 replicates), FASP-GPF (4 mass ranges, 4 replicates), FASP-GPF (8 mass ranges, 2 replicates), FASP-GPF (16 mass ranges,1 replicates), and FASP-HpH (8 fractions, 2 replicates) on linear ion trap mass spectrometer.

### 3.5 Proteins identified from tryptic digests on Q Exactive Orbitrap mass spectrometer

The results up to this point of this study indicated that the SDS-PAGE method with 16 fractions was the most advisable strategy for yeast protein identification. In order to quantify the effect of using a higher resolution mass spectrometer as opposed to a linear ion trap instrument, a second set of experiments were performed using a Q Exactive Orbitrap mass spectrometer to compare the SDS-PAGE method with 16 fractions, FASP-GPF (4 mass ranges, 4 replicates) and FASP-HpH fractionation (8 fractions, 2 replicates). As shown in Table 1, the FASP-GPF method allowed for the identification of 1035 proteins, the SDS-PAGE method produced 1357 protein identifications, and the FASP-HpH method generated 2134 identified proteins, with 859 proteins overlapped between all three methods (Figure 4). Combining the results collected from these three experiments on the Orbitrap instrument, there were a total of 2269 proteins found, with 94% of these proteins being identified by FASP-HpH method.

**Figure 4.**
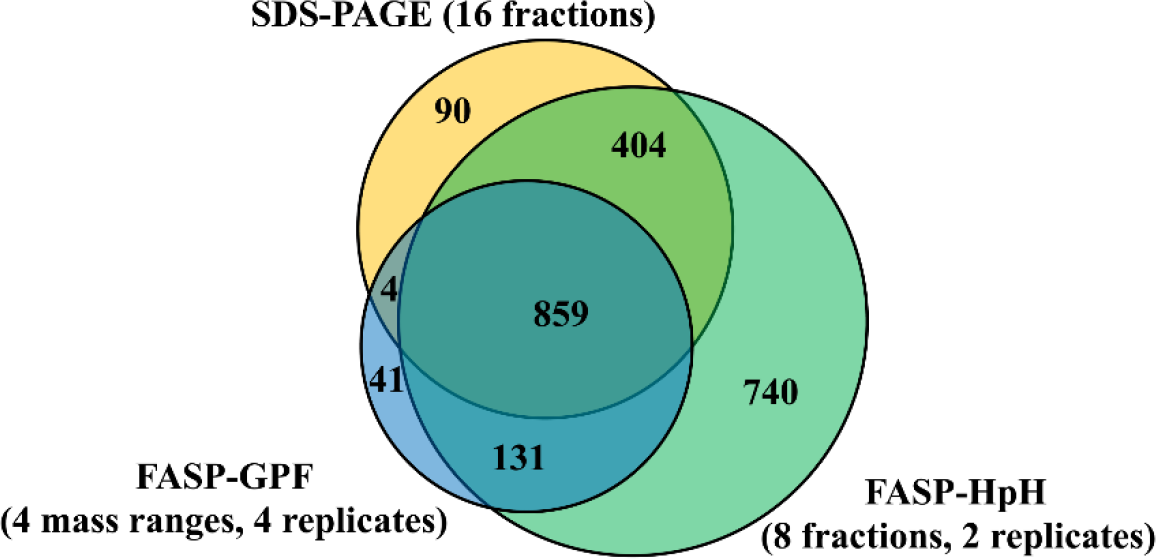
Venn diagram of proteins separately identified using SDS-PAGE (16 fractions), FASP-GPF (4 mass ranges, 4 replicates) and FASP-HpH (8 fractions, 2 replicates) on Q Exactive.

The peptides identified from these 859 overlapped proteins using three different proteomics approaches were analyzed further, considering both peptides and unique peptides, those which are found only by a particular method. Almost double the number of unique peptides and six times the number of total peptides were identified by FASP-HpH (8 fractions-2 replicates) (15963 unique peptides and 146635 total peptides) than by FASP-GPF (4 mass ranges, 4 replicates) (8458 unique peptides and 26950 total peptides). The SDS-PAGE method performed at an intermediate level, with 11791 unique and 105633 total peptides identified. The top 30 proteins, found in each of these three methods, ordered by sequence coverage, are presented in Table 2. There is a high degree of consistency between the methods, showing that the most abundant proteins are all identified in each experiment. The sequence coverage in percentage terms is generally higher using the FASP-HpH method, which correlates well with the fact that, on average, more peptides per proteins are identified using this approach on Q Exactive.

**Table 2.**
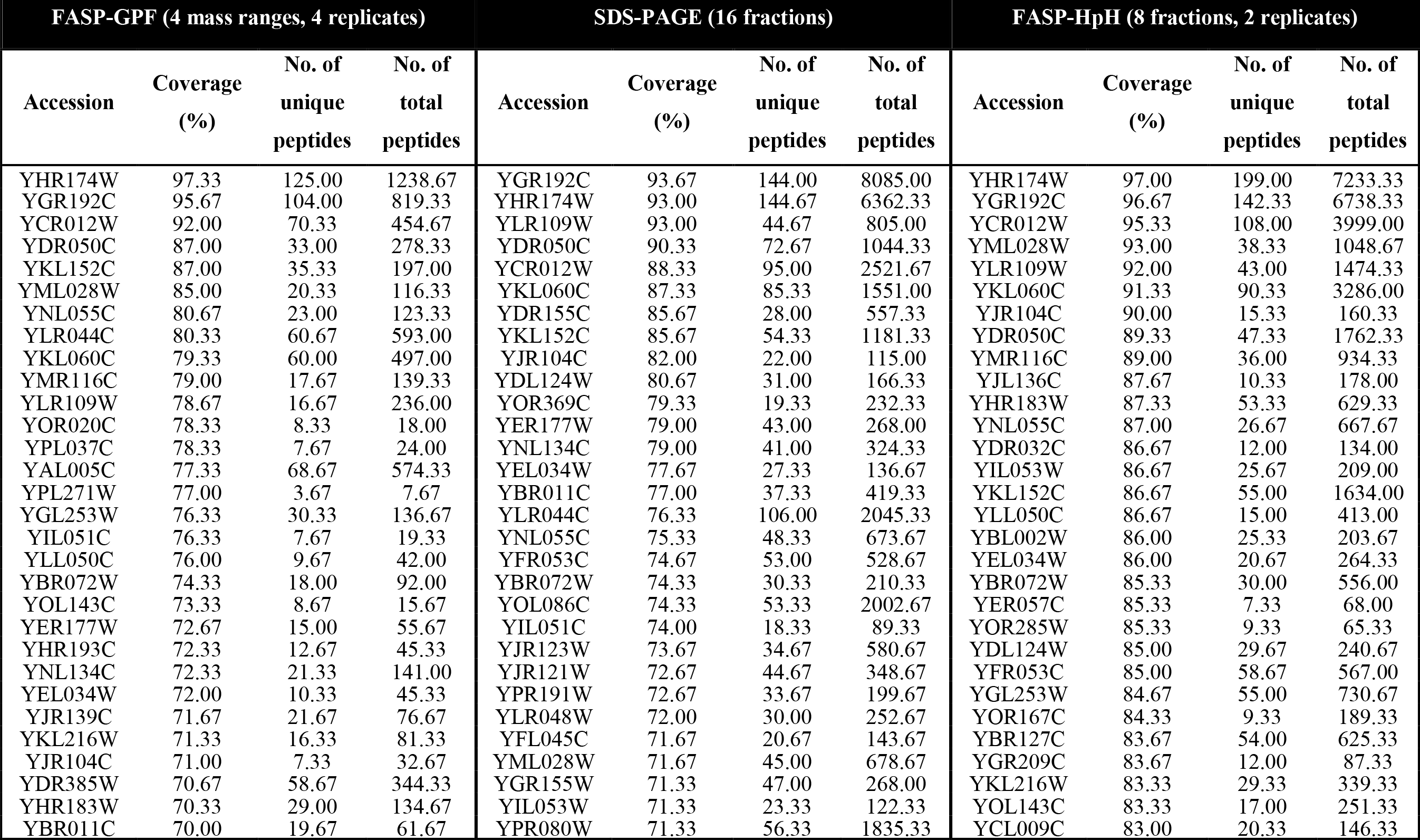
Top 30 proteins identified by FASP-GPF (4 mass ranges-4 replicates), SDS-PAGE (16 fractions) and FASP-HpH (8 fractions-2 replicates) experiments on Q Exactive Orbitrap mass spectrometer, ordered by sequence coverage.

The molecular weights of proteins separately identified by three methods on the Orbitrap instrument were analyzed. Most of the proteins were distributed between 20 kDa to 60 kDa, as shown in Figure 5A, with 665 proteins in FASP-GPF, 776 proteins in SDS-PAGE, and 1308 proteins in FASP-HpH. Similar results were found among the unique proteins separately identified using each of the three methods (Figure 5B). These distributions were similar to that of the identified proteins using a linear ion trap mass spectrometer (Figure 2).

**Figure 5.**
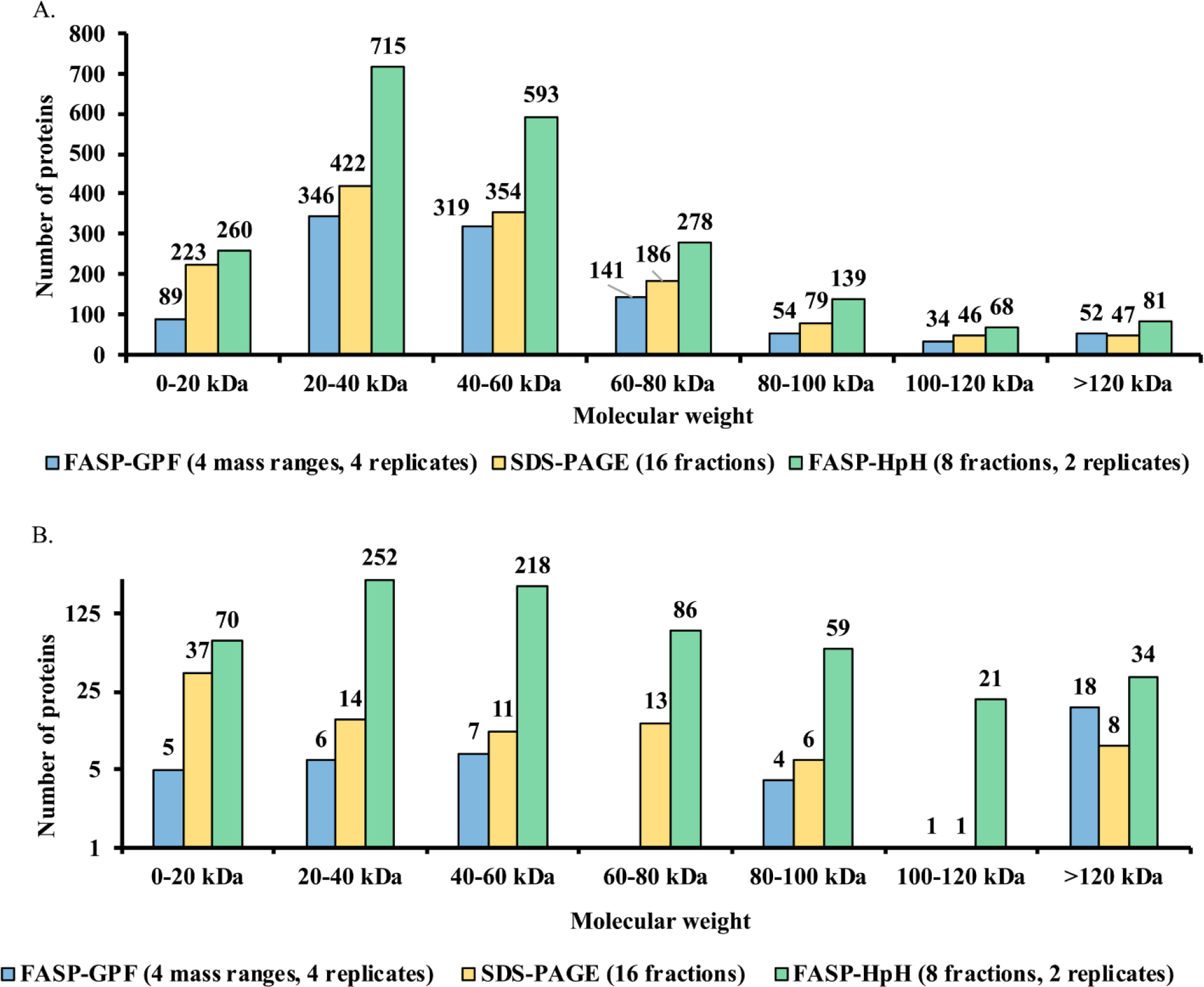
(A) Molecular weight distribution analysis of proteins separately identified using FASP-GPF (4 mass ranges, 4 replicates), SDS-PAGE (16 fractions) and FASP-HpH (8 fractions, 2 replicates) on Q Exactive. (B) Molecular weight distribution analysis of unique proteins separately identified using FASP-GPF (4 mass ranges, 4 replicates), SDS-PAGE (16 fractions) and FASP-HpH (8 fractions, 2 replicates) on Q Exactive.

### 3.6 mRNA abundance, Gene Ontology, and subcellular localization classification of proteins identified using an Orbitrap mass spectrometer

Proteins identified from SDS-PAGE, FASP-GPF and FASP-HpH experiments, using an Orbitrap mass spectrometer, were analyzed to compare the performance of each of these methods in identifying proteins when categorized by protein abundance, protein function, and protein subcellular localization.

In terms of protein abundance as evidenced by correlation with mRNA data, the FASP-HpH method was found to have a definite advantage in detecting proteins with lower abundance than the other two methods (Figure 6). At the two lowest abundance ranges, the FASP-HpH method identified almost twice as many proteins as the other approaches used.

**Figure 6.**
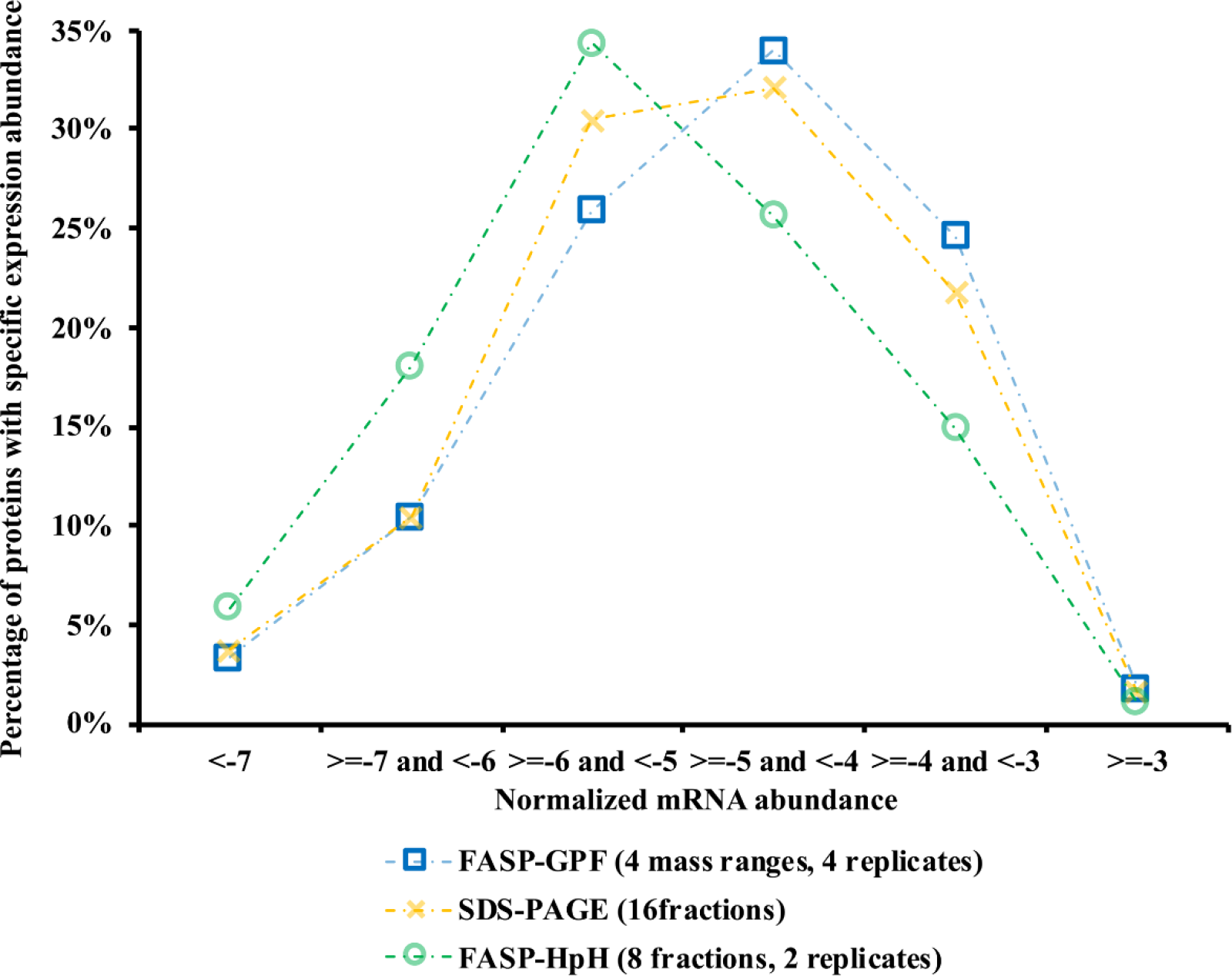
The correlation between protein abundance and mRNA expression analyzed using PARE, as described in methods. The X axis represents the normalized abundance of mRNAs which were categorized into six groups and used for indicating the theoretical value of corresponding protein expression. The minimum number indicates the category with the lowest abundance. The Y axis represents the percentage of proteins with specific expression abundance in all matched proteins to the mRNA reference in specific experiments, including FASP-GPF (4 mass ranges, 4 replicates), SDS-PAGE (16 fractions) and FASP-HpH (8 fractions, 2 replicates) on Q Exactive.

Gene ontology information for identified proteins was extracted from the Saccharomyces Genome Database using GO Slim mapper. These functional annotations revealed a broad distribution across a range of functions as shown in Figure 7. The three most abundant functional categories corresponded to proteins with function in ion binding (421 proteins from FASP-GPF, 483 proteins from SDS-PAGE, and 727 proteins from FASP-HpH), hydrolase activity (169 proteins from FASP-GPF, 206 proteins from SDS-PAGE, and 298 proteins from FASP-HpH), and transferase activity (172 proteins from FASP-GPF, 202 proteins from SDS-PAGE, and 326 proteins from FASP-HpH). Previous studies indicate that 70% of all enzymes bind metal ions since they are crucial for stabilizing protein structures and thereby affecting their function [43, 44]. Subcellular localization analysis of the combined set of identified proteins was performed based on the cellular component ontology within GO Slim mapper. The most common subcellular localizations associated with identified proteins were cytoplasm (56.9%), nucleus (42.1%), membrane (25.4%), and mitochondrion (24.7%) (Figure 8).

**Figure 7.**
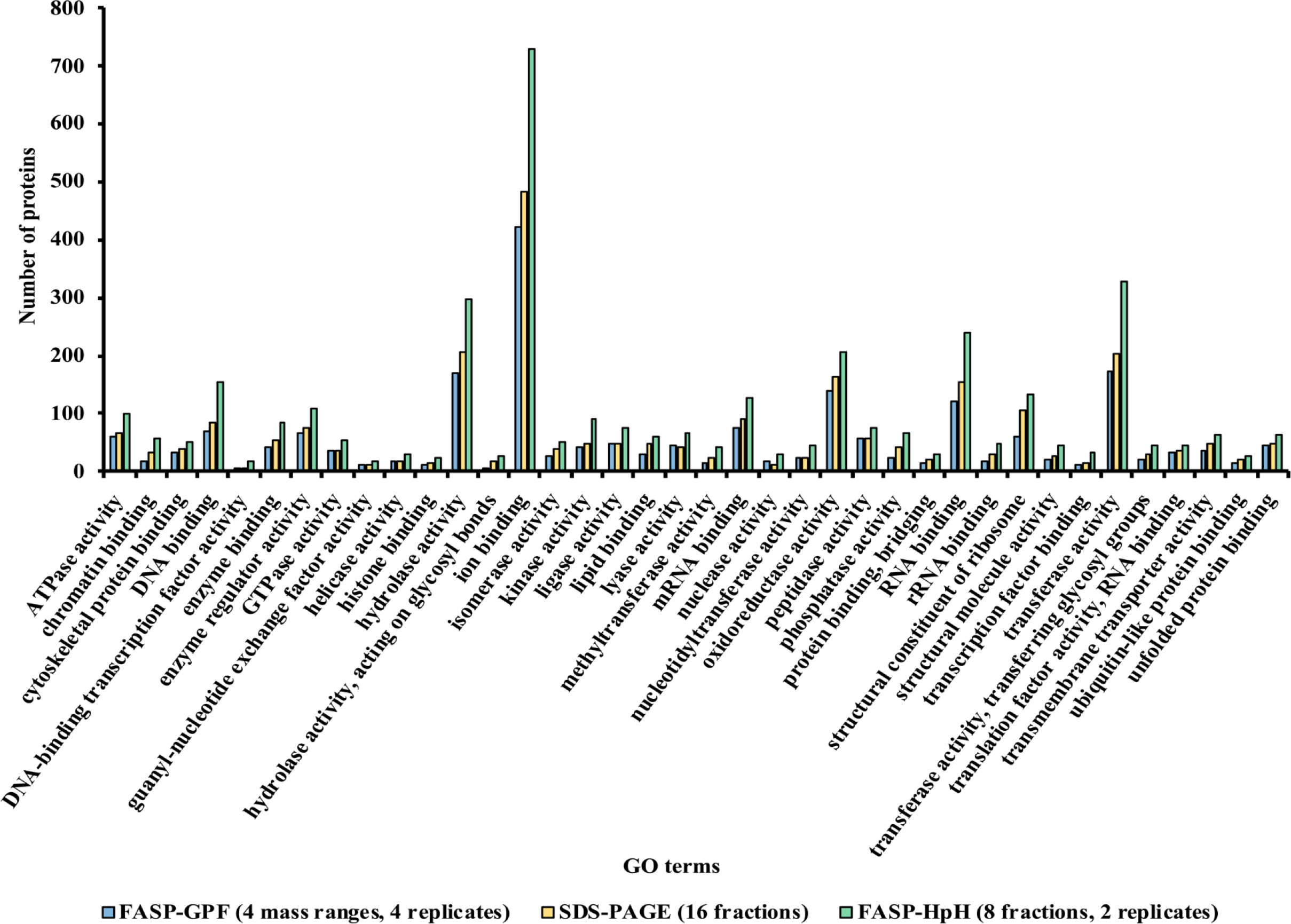
GO terms categorized with proteins separately identified using FASP-GPF (4 mass ranges-4 replicates), SDS-PAGE (16 fractions) and FASP-HpH (8 fractions, 2 replicates) according to their known functions based on gene ontology information in the Saccharomyces Genome Database using GO term mapper function.

**Figure 8.**
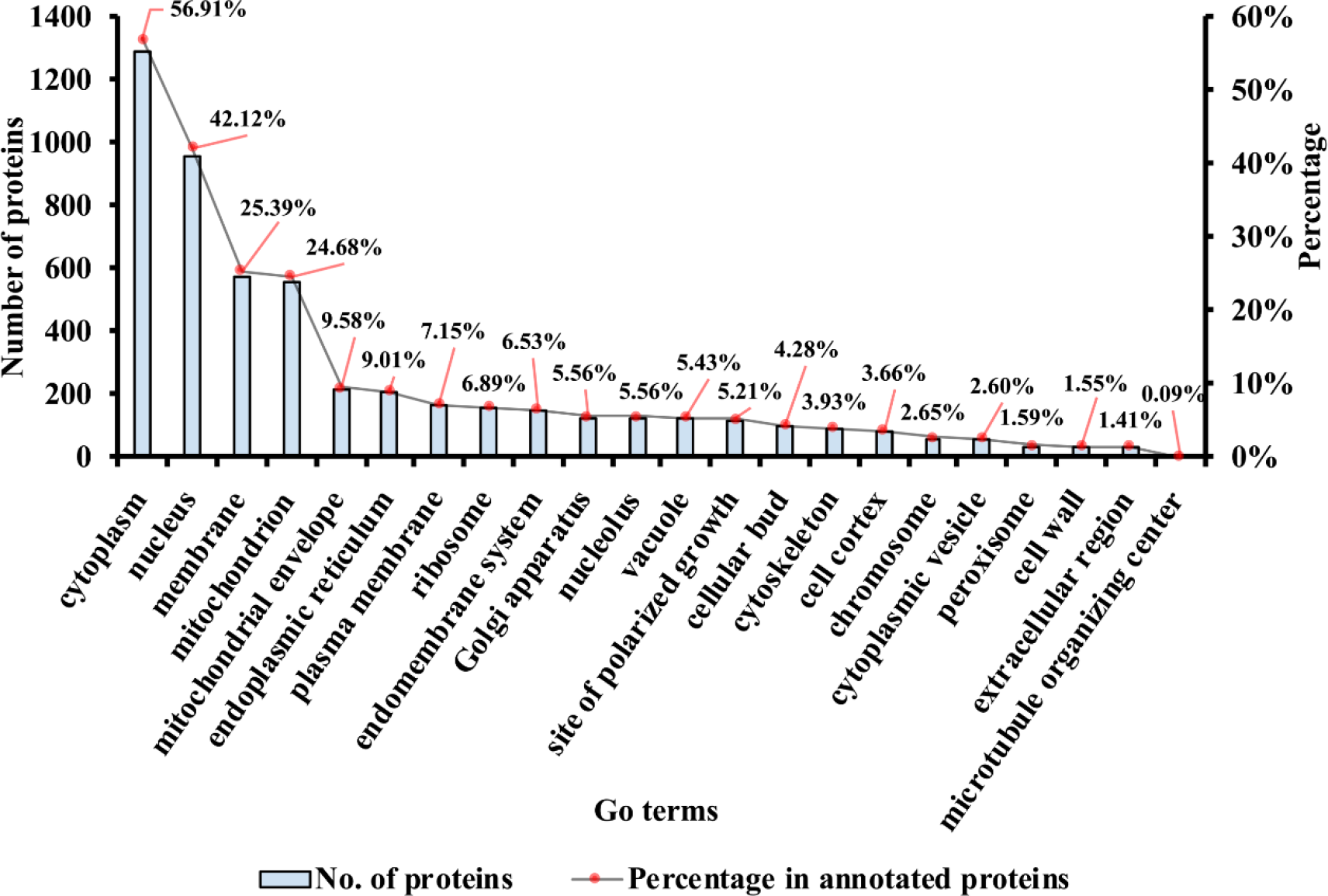
GO terms categorized with all combined proteins identified using FASP-GPF (4 mass ranges, 4 replicates), SDS-PAGE (16 fractions) and HpH (8 fractions, 2 replicates) on Q Exactive according to their cellular localization information in the Saccharomyces Genome Database using GO term mapper function.

### 3.7 Comparisons of fractionation approaches using two different mass spectrometers

It is possible to evaluate how much improvement in protein identification is made by using an Orbitrap mass spectrometer, rather than a linear ion trap mass spectrometer, using the data from those experiments performed with both: the SDS-PAGE (16 fractions), FASP-GPF (4 mass ranges, 4 replicates), and FASP-HpH (8 fractions, 2 replicates). The 16 fractions SDS-PAGE experiments showed that 12.5% more proteins were identified using a higher resolution instrument, with the number of reproducibly identified proteins increased from 1206 to 1357 (Table 1). However, the total number of peptides identified using the Orbitrap instrument was 176% higher than when using the linear ion trap (Table 1). For the FASP-GPF experiments (4 mass ranges, 4 replicates), using the higher resolution instrument increased the number of reproducibly identified proteins from 698 to 1035, an improvement of 48.3%. The number of peptides identified was increased by 83%. The greatest increase using the Q Exactive instrument was achieved with the FASP-HpH method, with a 210% increase in protein identification and an increase of 8.5 times in the number of peptides identified. More proteins with lower abundance were identified by Q Exactive (Figure 9), especially with FASP-GPF and FASP-HpH in the three lowest abundance categories. These results clearly illustrate the obvious advantages of using a higher resolution, higher speed instrument.

**Figure 9.**
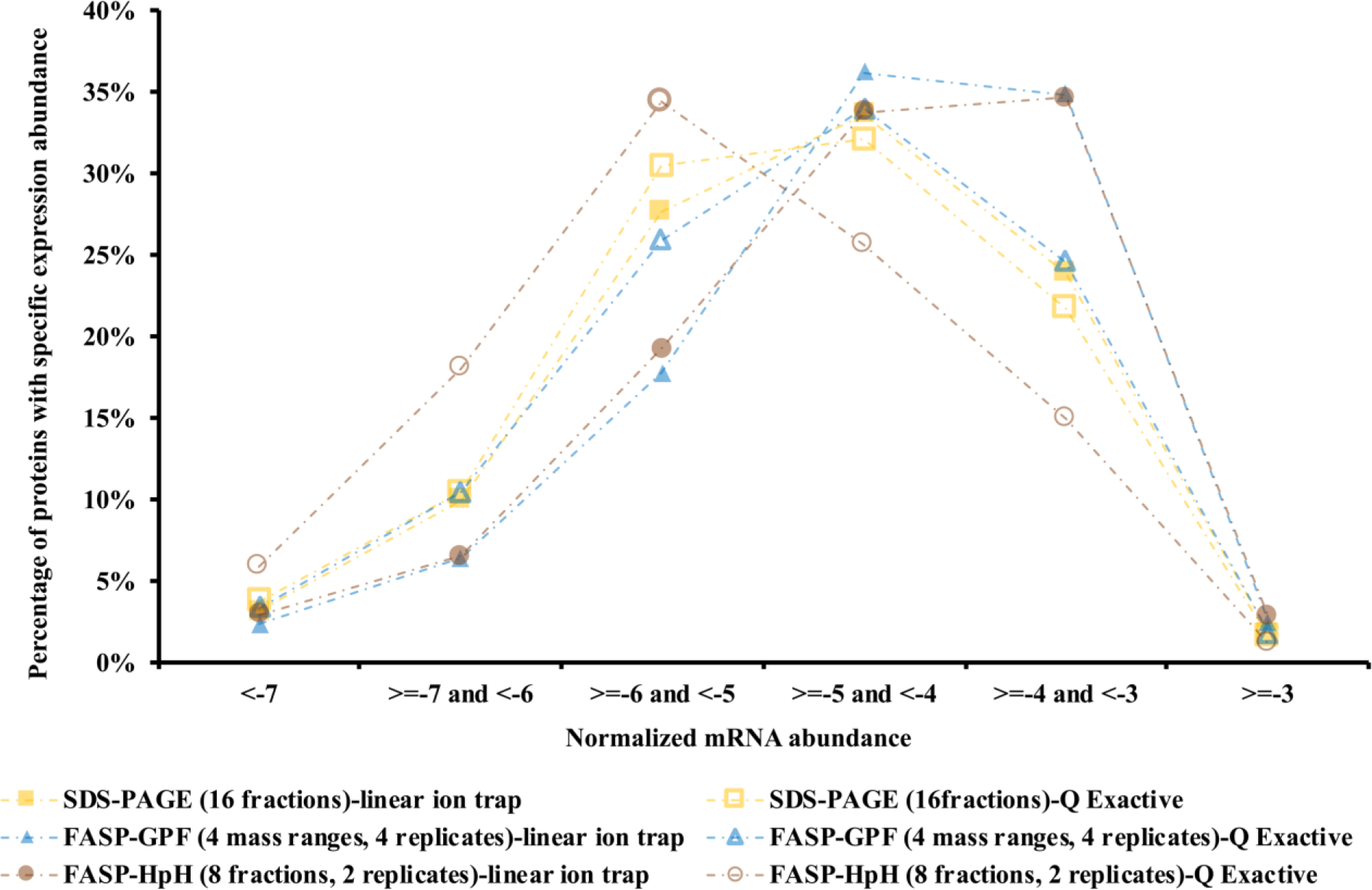
The correlation between protein abundance and mRNA expression analyzed using PARE, as described in methods. The X axis represents the normalized abundance of mRNAs which were categorized into six groups and used for indicating the theoretical value of corresponding protein expression. The minimum number indicates the category with the lowest abundance. The Y axis represents the percentage of proteins with specific expression abundance in all matched proteins to the mRNA reference in specific experiments, including FASP-GPF (4 mass ranges-4 replicates), SDS-PAGE (16 fractions) and FASP-HpH (8 fractions, 2 replicates) using Q Exactive and linear ion trap mass spectrometers, respectively.

## 4. DISCUSSION

Quantitative comparison of FASP-based in solution digestion and the traditional in-solution approach illustrated the FASP approach increased the number of identified proteins, which is consistent with previous reports [5, 21, 45]. Proteins are normally denatured before trypsin digestion to facilitate proteolysis, with highly concentrated urea commonly used during sample preparation due to its high efficiency in solubilization and denaturation of proteins. However, proteases are also subject to the denaturation effects of a high concentration of urea, which can interfere with enzymatic digestion, resulting in reduced proteolytic efficiency [21]. Therefore, complicated protein mixtures such as cellular lysates, which are usually solubilized in a high concentration of urea, are then diluted prior to protease digestion. However, a concentration of urea that still retains protease activity is often below the desirable concentration to fully denature the whole protein mixture, leading to sub-optimal digestion efficiency [46]. Additionally, urea is incompatible with mass spectrometric analysis, and therefore requires further desalting which may result in sample loss [47]. Many of these limitations are overcome by the relatively straightforward FASP sample preparation process [1]. The filter unit acts as a 'proteomic reactor' for detergent removal, buffer exchange, chemical modification and protein digestion [48]. Proteins are digested by trypsin in volatile ammonium bicarbonate buffer which can be removed in a vacuum centrifuge, producing a peptide mixture which is directly compatible with mass spectrometric analysis. FASP-based methods have been reported to improve throughput while reducing sample loss and increasing peptide recovery. Our results in this study show that the FASP digest approach does indeed lead to more proteins identified, and it is compatible with several different peptide fractionation strategies.

GPF has been shown in previous studies to be a highly effective fractionation technique when sufficient protein is available. However, the main drawback to this technique is that only a small portion of the whole sample is actually analyzed in the mass spectrometer, raising concerns regarding loss of information, especially in cases when there is only limited sample [20]. In GPF experiments, there is always a balance that must be achieved between reducing the survey scan mass range sufficiently to avoid under-sampling ions which may be present, and avoiding the potential loss of information caused by making the survey scan so small that most ions are excluded. Given a finite amount of starting material, analyzing more mass ranges with a smaller survey scan window means that a higher percentage of the sample is physically excluded from analysis in the mass spectrometer. This paradigm is one of the reasons why the ‘PAcIFIC’ data independent analysis technique, which can be thought of as an extreme version of gas phase fractionation, is typically performed on larger amounts of starting material [20, 49]. In this study, we found that using more injections of a smaller mass range in GPF experiments actually produced less protein identifications, and even in the best GPF experiments we performed, the results were still not as good as those obtained with other fractionation strategies. Interestingly, the Q Exactive Orbitrap instrument, with higher resolution, seems to compensate partly for the drawback of the GPF method, and reduced the difference between protein identification between FASP-GPF (4 bins-4 times) and SDS-PAGE (16 fractions) methods, indicating that the GPF method could be considered when it is coupled with a high resolution instrument with and the sample amount and experimental time are not limited.

Fractionation based on molecular weight using SDS-PAGE is one of the most common pre-fractionation methods currently in use, due to its robustness and effective performance in achieving maximal proteome coverage [12]. Although more manual handling is involved when using SDS-PAGE fractionation, gels are also useful for separating complex proteins and facilitating the removal of contaminants that may interfere with mass spectrometry analysis [1]. Another obvious advantage of SDS-PAGE is that this procedure allows practically any protein samples to be visualized and studied because SDS is a highly effective protein solubilization agent.

While the most important consideration during proteomics analysis is usually the yield of proteins and peptides, the time required for analysis is also an important consideration. Cutting more gel slices and recovering peptides from gel pieces are time-consuming experiments, and mass spectrometer instrument time can also be a significant concern. In this study, the SDS-PAGE gel slice fractionation method proved to be an effective and consistent sample preparation method. The fact that only a marginal increase in protein identifications occurred when the number of gel slices used was increased from 16 to 32 is an interesting finding. This indicates that fractionation into 16 gel slices was sufficient to reduce the complexity of this peptide mixture so that it was compatible with the analytical capacity of the lower resolution linear ion trap mass spectrometer. This is in contrast to previous studies which reported a linear relationship between the number of gel slices and the number of proteins identified [50]. This finding reinforces the idea that there is no ideal number of gel slices that can be applied to every experiment, because it is highly dependent on both sample type and instrumental performance, and it needs to be empirically determined.

High pH reversed phase fractionation, as a first-dimension peptide separation technique, performs exceptionally well in peptide separation and is highly orthogonal to low pH reversed phase separation used for MS analysis [25]. This efficiency of peptide fractionation can increase subsequent protein identifications and sequence coverage, which helps achieve deeper proteome coverage. In this study, we used a FASP-HpH method which combines the advantages of FASP in solution digestion which improved the peptide recovery, and HpH fractionation which increased the fractionation efficiency. The application of this method using a high resolution instrument improved both peptide and protein identification when compared with GPF and SDS-PAGE approaches. In a previous report, a comparison was performed between HpH fractionation and a peptide isoelectric focusing approach, which also indicated better performance of HpH in terms of proteome coverage [51, 52]. In our hands, FASP-HpH represents a relatively straightforward proteomics workflow, and when using a linear ion trap instrument produced results comparable to FASP-GPF, but was less effective than SDS-PAGE. However, when using a faster, high resolution Orbitrap mass spectrometer, the FASP-HpH approach was clearly the best approach to use, and was able to achieve a remarkable depth of proteome coverage.

A proteome is a collection of gene products expressed at both low and high levels. Effective identification and quantitative comparison of low-abundance proteins is still a tremendous challenge in proteomic analysis [53]. It is worth mentioning that many biologically relevant changes in the proteome occur at the middle to low range of the protein abundance scale. Ideally, shotgun proteomics would facilitate the identification of an entire proteome with 100% protein sequence coverage. In reality, the large dynamic range and complexity of cellular proteomes result in oversampling of abundant proteins, while peptides from low abundance proteins are under-sampled or remain undetected [53]. Protein levels are linked to mRNA expression by the process of translation, and mRNA expression data have been widely used to directly predict or model protein expression [54–57]. Although the mRNA level of a gene is not always sufficient by itself to predict the expression level of a particular protein due to post-transcriptional and translation regulation [58, 59], transcriptome data unquestionably provides an excellent framework for prediction of protein abundances, especially in large data sets. The correlation analysis of both transcription and translation levels has been applied in a range of biological contexts to provide insights into the study of many molecular mechanisms [60–62]. In our study, correlation analysis between protein and mRNA abundance revealed that fractionation using FASP-HpH and nano LC-MS/MS with a high resolution mass spectrometer performed much better than the other methods examined in identifying less abundant proteins, which may be partly attributable to the enhanced separation capacity of this method, and faster scan speed of the instrument, leading to identification of more peptides.

## 5. CONCLUSIONS

The information produced in this study provides a valuable method optimization framework for researchers developing proteomics analysis projects. Although the FASP-HpH method contributed most of the total protein identifications from the experiments performed using an Orbitrap mass spectrometer, different approaches should always be considered when undertaking proteomics experiments. For example, although in our hands the GPF method did not perform particularly well in identifying proteins from yeast, in a previous study we reported it to have better performance in identifying more reproducible proteins in grapevines, when compared to SDS-PAGE fractionation [19]. The GPF method is cost neutral and can still be of utility as a complementary method in proteomics analysis. In addition to the methods applied in this study, there are many other tools and techniques available for identification and characterization of complex protein mixtures, all of which have different strengths and limitations. Although it is common to utilize more than one method to achieve more protein identifications [8, 20], the sample preparation and fractionation strategies used for shotgun proteomics analysis are always sample-dependent. In this study, SDS-PAGE, as the most common sample preparation method for any kind of protein samples, showed good performance on both low- and high-resolution instruments, which makes it optimal for many research groups. FASP-HpH was the best available method for shotgun proteomics of the yeast samples when applied on a faster, higher resolution instrument as it considerably increased the scope and scale of peptide and protein identification.

## ACKNOWLEDGEMENTS

The authors wish to thank Hugh Goold and Ian Paulsen for providing yeast cell culture starting materials. LD would like to acknowledge scholarship support from the China scholarship Council This work was supported by Macquarie University, and aspects of this research were conducted at the Australian Proteome Analysis Facility.

## REFERENCES

[1] P. Feist, A.B. Hummon, Proteomic challenges: sample preparation techniques for microgram-quantity protein analysis from biological samples, International Journal of Molecular Sciences, 16 (2015), pp. 3537–3563.

[2] J. Berg, J. Tymoczko, L. Stryer, Protein structure and function, Biochemistry, W H Freeman, New York, (2002), pp. 159–173.

[3] Y.Y. Zhang, B.R. Fonslow, B. Shan, M.C. Baek, J.R. Yates, Protein analysis by shotgun/bottom-up proteomics, Chemical Reviews, 113 (2013), pp. 2343–2394.

[4] T. Léger, C. Garcia, M. Videlier, J.-M. Camadro, Label-free quantitative proteomics in yeast, in: F. Devaux (Ed.) Yeast Functional Genomics, Humana Press, (2016), pp. 289–307.

[5] K.R. Ludwig, M.M. Schroll, A.B. Hummon, Comparison of in-solution, FASP, and S-trap based digestion methods for bottom-up proteomic studies, Journal of Proteome Research, 17 (2018), pp. 2480–2490.

[6] L.-R. Yu, N. A. Stewart, T. D. Veenstra, Proteomics: the deciphering of the functional genome, in: G.S. Ginsburg, H.F. Willard (Eds.) Essentials of Genomic and Personalized Medicine, Academic Press, (2010), pp. 89–96.

[7] H.M. Mottaz-Brewer, A.D. Norbeck, J.N. Adkins, N.P. Manes, C. Ansong, L. Shi, Y. Rikihisa, T. Kikuchi, S.W. Wong, R.D. Estep, F. Heffron, L. Pasa-Tolic, R.D. Smith, Optimization of proteomic sample preparation procedures for comprehensive protein characterization of pathogenic systems, Journal of Biomolecular Techniques, 19 (2008), pp. 285–295.

[8] M. Mirzaei, K. Pushpitha, L. Deng, N. Chitranshi, V. Gupta, R. Rajput, A.B. Mangani, Y. Dheer, A. Godinez, M.J. McKay, K. Kamath, D. Pascovici, J.X. Wu, G.H. Salekdeh, T. Karl, P.A. Haynes, S.L. Graham, V.K.J.M.N. Gupta, Upregulation of proteolytic pathways and altered protein biosynthesis underlie retinal pathology in a mouse model of Alzheimer’s disease, Molecular Neurobiology, 56 (2019), pp. 6017–6034.

[9] A. Bodzon-Kulakowska, A. Bierczynska-Krzysik, T. Dylag, A. Drabik, P. Suder, M. Noga, J. Jarzebinska, J. Silberring, Methods for samples preparation in proteomic research, Journal of Chromatography B, 849 (2007), pp. 1–31.

[10] F.E. Ahmed, Sample preparation and fractionation for proteome analysis and cancer biomarker discovery by mass spectrometry, Journal of Separation Science, 32 (2009), pp. 771–798.

[11] S.S. Thakur, T. Geiger, B. Chatterjee, P. Bandilla, F. Frohlich, J. Cox, M. Mann, Deep and highly sensitive proteome coverage by LC-MS/MS without prefractionation, Molecular & Cellular Proteomics, 10 (2011).

[12] Y. Fang, D.P. Robinson, L.J. Foster, Quantitative analysis of proteome coverage and recovery rates for upstream fractionation methods in proteomics, Journal of Proteome Research, 9 (2010), pp. 1902–1912.

[13] K. Chandramouli, P.Y. Qian, Proteomics: challenges, techniques and possibilities to overcome biological sample complexity, Human Genomics and Proteomics, 2009 (2009).

[14] U. Hellman, Sample preparation by SDS/PAGE and in-gel digestion, Proteomics in Functional Genomics, 88 (2000), pp. 43–54.

[15] B. Granvogl, M. Ploscher, L.A. Eichacker, Sample preparation by in-gel digestion for mass spectrometry-based proteomics, Analytical and Bioanalytical Chemistry 389 (2007), pp. 991–1002.

[16] Y.V. Karpievitch, A.D. Polpitiya, G.A. Anderson, R.D. Smith, A.R. Dabney, Liquid chromatography mass spectrometry-based proteomics: biological and technological aspects, The Annals of Applied Statistics 4(2010), pp. 1797–1823.

[17] Z.J. Cao, H.Y. Tang, H. Wang, Q. Liu, D.W. Speicher, Systematic comparison of fractionation methods for in-depth analysis of plasma proteomes, Journal of Proteome Research, 11 (2012), pp. 3090–3100.

[18] I.R. Leon, V. Schwammle, O.N. Jensen, R.R. Sprenger, Quantitative assessment of in-solution digestion efficiency identifies optimal protocols for unbiased protein analysis, Molecular & Cellular Proteomics, 12 (2013), pp. 2992–3005.

[19] I.S. George, A.Y. Fennell, P.A. Haynes, Protein identification and quantification from riverbank grape, Vitis riparia: comparing SDS-PAGE and FASP-GPF techniques for shotgun proteomic analysis, Proteomics, 15 (2015), pp. 3061–3065.

[20] L. Breci, E. Hattrup, M. Keeler, J. Letarte, R. Johnson, P.A. Haynes, Comprehensive proteornics in yeast using chromatographic fractionation, gas phase fractionation, protein gel electrophoresis, and isoelectric focusing, Proteomics, 5 (2005), pp. 2018–2028.

[21] L.A. Weston, K.M. Bauer, A.B. Hummon, Comparison of bottom-up proteomic approaches for LC-MS analysis of complex proteomes, Analytical Methods, 5 (2013), pp. 4615–4621.

[22] A. Scherl, S.A. Shaffer, G.K. Taylor, H.D. Kulasekara, S.I. Miller, D.R. Goodlett, Genome-specific gas-phase fractionation strategy for improved shotgun proteomic profiling of proteotypic peptides, Analytical Chemistry, 80 (2008), pp. 1182–1191.

[23] J. Kennedy, E.C. Yi, Use of gas-phase fractionation to increase protein identifications : application to the peroxisome, in: D. Pflieger, J. Rossier (Eds.) Organelle Proteomics, Humana Press, (2008), pp. 217–228.

[24] O. Coleman, M. Henry, M. Clynes, P. Meleady, Filter-aided sample preparation (FASP) for improved proteome analysis of recombinant Chinese hamster ovary cells, in: P. Meleady (Ed.) Heterologous Protein Production in CHO Cells, Humana Press, (2017), pp. 187–194.

[25] T.S. Batth, C. Francavilla, J.V. Olsen, Off-line high-pH reversed-phase fractionation for in-depth phosphoproteomics, Journal of Proteome Research, 13 (2014), pp. 6176–6186.

[26] F. Yang, Y.F. Shen, D.G. Camp, R.D. Smith, High-pH reversed-phase chromatography with fraction concatenation for 2D proteomic analysis, Expert Review of Proteomics, 9 (2012), pp. 129–134.

[27] T.T. Zhang, J. Lei, H.J. Yang, K. Xu, R. Wang, Z.Y. Zhang, An improved method for whole protein extraction from yeast Saccharomyces cerevisiae, Yeast, 28 (2011), pp. 795–798.

[28] D. Wessel, U.I. Flugge, A method for the quantitative recovery of protein in dilute solution in the presence of detergents and lipids, Analytical Biochemistry, 138 (1984), pp. 141–143.

[29] V. Vaibhav, E.L. Thompson, D.A. Raftos, P.A. Haynes, Potential protein biomarkers of QX disease resistance in selectively bred Sydney Rock Oysters, Aquaculture, 495 (2018), pp. 144–152.

[30] S. Rattanakan, I. George, P.A. Haynes, G.R. Cramer, Relative quantification of phosphoproteomic changes in grapevine (Vitis vinifera L.) leaves in response to abscisic acid, Horticulture Research, 3 (2016).

[31] Y.Q. Wu, M. Mirzaei, D. Pascovici, J.M. Chick, B.J. Atwell, P.A. Haynes, Quantitative proteomic analysis of two different rice varieties reveals that drought tolerance is correlated with reduced abundance of photosynthetic machinery and increased abundance of ClpD1 protease, Journal of Proteomics, 143 (2016), pp. 73–82.

[32] R. Craig, R.C. Beavis, A method for reducing the time required to match protein sequences with tandem mass spectra, Rapid Communications in Mass Spectrometry, 17 (2003), pp. 2310–2316.

[33] R.C. Beavis, Using the global proteome machine for protein identification, New and Emerging Proteomic Techniques, 328 (2006), pp. 217–228.

[34] K.A. Neilson, T. Keighley, D. Pascovici, B. Cooke, P.A. Haynes, Label-free quantitative shotgun proteomics using normalized spectral abundance factors, in: M. Zhou, T. Veenstra (Eds.) Proteomics for Biomarker Discovery, Humana Press, (2013), pp. 205–222.

[35] D.C. Handler, P.A. Haynes, An experimentally-derived measure of inter-replicate variation in reference samples: the same-same permutation methodology, bioRxiv, (2019).

[36] M. Mirzaei, D. Pascovici, J.X. Wu, J. Chick, Y.Q. Wu, B. Cooke, P. Haynes, M.P. Molloy, TMT one-stop shop: from reliable sample preparation to computational analysis platform, in: S. Keerthikumar, S. Mathivanan (Eds.) Proteome Bioinformatics, (2017), pp. 45–66.

[37] A. Fathi, M. Mirzaei, B. Dolatyar, M. Sharifitabar, M. Bayat, E. Shahbazi, J. Lee, M. Javan, S.C. Zhang, V. Gupta, B. Lee, P.A. Haynes, H. Baharvand, G.H. Salekdeh, Discovery of novel cell surface markers for purification of embryonic dopamine progenitors for transplantation in Parkinson’s disease animal models, Molecular & Cellular Proteomics, 17 (2018), pp. 1670–1684.

[38] M. Mirzaei, N. Soltani, E. Sarhadi, I.S. George, K.A. Neison, D. Pascovici, S. Shahbazian, P.A. Haynes, B.J. Atwell, G.H. Salekdeh, Manipulating root water supply elicits major shifts in the shoot proteome, Journal of Proteome Research, 13 (2014), pp. 517–526.

[39] J.M. Cherry, E.L. Hong, C. Amundsen, R. Balakrishnan, G. Binkley, E.T. Chan, K.R. Christie, M.C. Costanzo, S.S. Dwight, S.R. Engel, D.G. Fisk, J.E. Hirschman, B.C. Hitz, K. Karra, C.J. Krieger, S.R. Miyasato, R.S. Nash, J. Park, M.S. Skrzypek, M. Simison, S. Weng, E.D. Wong, Saccharomyces Genome Database: the genomics resource of budding yeast, Nucleic Acids Research, 40 (2012), pp. D700–D705.

[40] M.S. Skrzypek, R.S. Nash, E.D. Wong, K.A. MacPherson, S.T. Hellerstedt, S.R. Engel, K. Karra, S. Weng, T.K. Sheppard, G. Binkley, M. Simison, S.R. Miyasato, J.M. Cherry, Saccharomyces genome database informs human biology, Nucleic Acids Research, 46 (2018), pp. D736–D742.

[41] E.Z. Yu, A.E. Burba, M. Gerstein, PARE: a tool for comparing protein abundance and mRNA expression data, BMC Bioinformatics, 8 (2007).

[42] Y.L. Lee, C.K. Lee, Transcriptional response according to strength of calorie restriction in Saccharomyces cerevisiae, Molecules and Cells, 26 (2008), pp. 299–307.

[43] M. Petukh, E. Alexov, Ion binding to biological macromolecules, Asian Journal of Physics, 23 (2014), pp. 735–744.

[44] C.H. Lu, Y.F. Lin, J.J. Lin, C.S. Yu, Prediction of metal ion–binding sites in proteins using the fragment transformation method, Plos One, 7 (2012).

[45] W.Q. Wang, O.N. Jensen, I.M. Moller, K.H. Hebelstrup, A. Rogowska-Wrzesinska, Evaluation of sample preparation methods for mass spectrometry-based proteomic analysis of barley leaves, Plant Methods, 14 (2018).

[46] E.I. Chen, D. Cociorva, J.L. Norris, J.R. Yates, Optimization of mass spectrometry-compatible surfactants for shotgun proteomics, Journal of Proteome Research, 6 (2007), pp. 2529–2538.

[47] S.C. Kim, Y. Chen, S. Mirza, Y.D. Xu, J. Lee, P.S. Liu, Y.M. Zhao, A clean, more efficient method for in-solution digestion of protein mixtures without detergent or urea, Journal of Proteome Research, 5 (2006), pp. 3446–3452.

[48] J.R. Wisniewski, A. Zougman, N. Nagaraj, M. Mann, Universal sample preparation method for proteome analysis, Nature Methods, 6 (2009), pp. 359–362.

[49] A. Panchaud, A. Scherl, S.A. Shaffer, P.D. von Haller, H.D. Kulasekara, S.I. Miller, D.R. Goodlett, Precursor acquisition independent from ion count: how to dive deeper into the proteomics ocean, Analytical Chemistry, 81 (2009), pp. 6481–6488.

[50] A. Shevchenko, H. Tomas, J. Havlis, J.V. Olsen, M. Mann, In-gel digestion for mass spectrometric characterization of proteins and proteomes, Nature Protocols, 1 (2006), pp. 2856–2860.

[51] H. Wang, S. Sun, Y. Zhang, S. Chen, P. Liu, B. Liu, An off-line high pH reversed-phase fractionation and nano-liquid chromatography-mass spectrometry method for global proteomic profiling of cell lines, Journal of Chromatography B, 974 (2015), pp. 90–95.

[52] D.R. Stein, X.J. Hu, S.J. McCorrister, G.R. Westmacott, F.A. Plummer, T.B. Ball, M.S. Carpenter, High pH reversed-phase chromatography as a superior fractionation scheme compared to off-gel isoelectric focusing for complex proteome analysis, Proteomics, 13 (2013), pp. 2956–2966.

[53] B.R. Fonslow, P.C. Carvalho, K. Academia, S. Freeby, T. Xu, A. Nakorchevsky, A. Paulus, J.R. Yates, Improvements in proteomic metrics of low abundance proteins through proteome equalization using ProteoMiner prior to MudPIT, Journal of Proteome Research, 10 (2011), pp. 3690–3700.

[54] L. Nie, G. Wu, W. Zhang, Correlation of mRNA expression and protein abundance affected by multiple sequence features related to translational efficiency in Desulfovibrio vulgaris: a quantitative analysis, Genetics, 174 (2006), pp. 2229–2243.

[55] L. Ponnala, Y. Wang, Q. Sun, K.J. van Wijk, Correlation of mRNA and protein abundance in the developing maize leaf, The Plant Journal, 78 (2014), pp. 424–440.

[56] D. Greenbaum, C. Colangelo, K. Williams, M. Gerstein, Comparing protein abundance and mRNA expression levels on a genomic scale, Genome Biology, 4 (2003).

[57] A.M. Mehdi, R. Patrick, T.L. Bailey, M. Boden, Predicting the dynamics of protein abundance, Molecular & Cellular Proteomics, 13 (2014), pp. 1330–1340.

[58] J.M. Laurent, C. Vogel, T. Kwon, S.A. Craig, D.R. Boutz, H.K. Huse, K. Nozue, H. Walia, M. Whiteley, P.C. Ronald, Protein abundances are more conserved than mRNA abundances across diverse taxa, Proteomics, 10 (2010), pp. 4209–4212.

[59] Y. Liu, A. Beyer, R. Aebersold, On the dependency of cellular protein levels on mRNA abundance, Cell, 165 (2016), pp. 535–550.

[60] A. Ghazalpour, B. Bennett, V.A. Petyuk, L. Orozco, R. Hagopian, I.N. Mungrue, C.R. Farber, J. Sinsheimer, H.M. Kang, N. Furlotte, Comparative analysis of proteome and transcriptome variation in mouse, Plos Genetics, 7 (2011).

[61] D. Garcia-Seco, M. Chiapello, M. Bracale, C. Pesce, P. Bagnaresi, E. Dubois, L. Moulin, C. Vannini, R. Koebnik, Transcriptome and proteome analysis reveal new insight into proximal and distal responses of wheat to foliar infection by Xanthomonas translucens, Scientific Reports, 7 (2017).

[62] C. Manzoni, D.A. Kia, J. Vandrovcova, J. Hardy, N.W. Wood, P.A. Lewis, R. Ferrari, Genome, transcriptome and proteome: the rise of omics data and their integration in biomedical sciences, Briefings in Bioinformatics 19 (2016), pp. 286–302.

